# Evaluation of Nanopore direct RNA sequencing updates for modification detection

**DOI:** 10.1101/2025.05.01.651717

**Authors:** Neda Ghohabi Esfahani, Andrew J Stein, Stuart Akeson, Talia Tzadikario, Miten Jain

## Abstract

Nanopore technology can directly sequence intact RNA molecules, offering a unique capability to read native modifications. Oxford Nanopore Technologies recently updated its direct RNA sequencing technology from RNA002 to RNA004 chemistry. This update included an improved basecaller (Dorado) for increased sequencing accuracy, and ionic current models for de novo detection of four RNA modifications. Using a single RNA extraction from GM12878 B-lymphocyte cell line, we compared RNA002 and RNA004 sequencing chemistries and evaluated Dorado modification calling accuracy. We computed U-to-C mismatches, previously used to identify putative pseudouridine sites, and ran m6anet for identifying putative N6-methyladenosine sites. Dorado results for each respective modification showed both global and site-specific differences when compared to RNA002 results. We used DRS data from in vitro transcription of GM12878 genomic DNA as well as synthetic oligonucleotides to evaluate Dorado modification calling performance. Dorado’s pseudouridine model achieved 96–98% for both accuracy and F1-score. Similarly, Dorado’s N6-methyladenosine model achieved 94–98% accuracy, 96–99% F1-score. Our results demonstrate that Nanopore Direct RNA sequencing could simultaneously detect pseudouridine, N6-methyladenosine, 5-methylcytosine, and inosine modifications on individual mRNA strands. It is critical to validate these calls using orthogonal methods (e.g., Liquid Chromatography with Tandem Mass Spectrometry) for increased confidence.

**Graphical Abstract:** 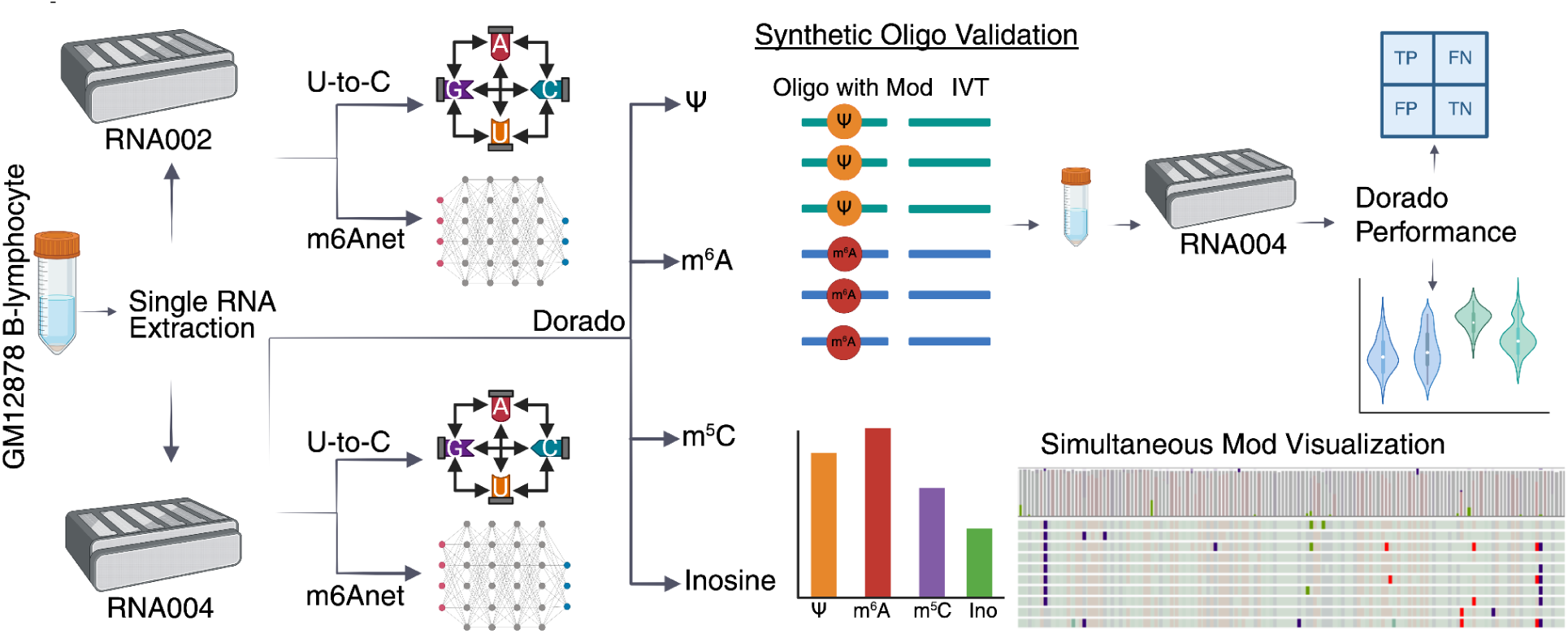

## Introduction

Nanopore direct RNA sequencing (DRS) is the only technology that allows for the long readouts of native RNA molecules^1,2^. Briefly, DRS involves the translocation of an RNA strand through a biological nanopore, where the ionic current is influenced by the nucleotide composition in the nanopore vestibule as they pass through. These fluctuations in ionic current are then interpreted (basecalled) into RNA sequences by deep learning algorithms. An advantage of DRS is that the native RNA molecules are measured, rather than a synthetic DNA copy^2^. Another key feature of DRS is the direct, simultaneous detection of RNA modifications at a single-molecule level^1,3–6^. RNA modifications are chemical alterations to ribonucleotides, with over 170 unique types currently identified^7^. These modifications alter the behavior across diverse classes of RNA, impacting translation^8^, stability^9^, and molecular structure^10^, with more roles likely to be uncovered with continued research^11^.

As with all models, the basecaller can make errors. Nucleotide modifications have been identified as one potential source of these errors. For example, uridine modifications were semi-systematically miscalled as a cytosine in data generated using RNA002 chemistry and basecalled using Guppy^5,6^. Alternatively, ionic current models were developed to interpret specific modifications. For example, m6anet^12^ differentiated N6-methyladenosine (m^6^A) from adenosine using ionic current and basecalled sequence data. Both the miscall error techniques and ionic current models for modification calling required continued development as DRS technology progressed.

The majority of studies utilizing DRS used RNA002 sequencing chemistry, which was available from 2019 to 2024. In 2024, ONT updated its DRS sequencing chemistry to RNA004^13^. This update in sequencing chemistry used a new RNA-specific Nanopore, requiring new basecalling models. This meant that the traditional RNA modification identification techniques from RNA002 were no longer directly applicable, and the strategies required validation before being applied to RNA004 data. Following RNA004, ONT released an updated basecaller Dorado^14^, with RNA modification calling capability^15^ for pseudouridine (Ψ), m^6^A, 5-methylcytosine (m^5^C), and inosine. None of these modification callers were supported with independent verification by the research community at the time of their release. Orthogonal validation techniques are crucial as modification-calling strategies advance. Validation can come in the form of true negative *in vitro* transcription (IVT) derived RNA^4,16^, enzymatic knockdowns and knockouts^17,18^, liquid chromatography with tandem mass spectrometry (LC-MS/MS)^17,19^, and sequencing of synthetic oligonucleotides with and without modifications^20^.

In this work, we compared RNA002 against RNA004 sequencing chemistry. We used a single extraction of RNA and sequenced it using both chemistries, allowing us to assume a constant baseline. Our analysis focused on the changes in alignment identity, insertion and deletion rates, and base substitution rates for these data. We also compared RNA004 modification calling methods (Dorado-based) with RNA002 modification calling methods (U-to-C mismatches, m6anet). To minimize false positives, we computed an IVT-controlled modification occupancy rate for each position, producing a subset of higher confidence putative modification sites^21^. For a subset of sequence contexts, we analyzed synthetic oligonucleotides to approximate a “known” modification state for Ψ and m^6^A. These synthetic controls provided validation for a set of modification calls by the novel RNA004-based modification callers.

## Results

### RNA004 chemistry has improved sequencing performance compared with RNA002 chemistry

The RNA002 sequencing experiments from Workman et al. had yielded 9,729,412 reads with a read N50 of 1.36kb. The RNA004 sequencing experiment produced 14,494,046 reads with a read N50 of 1.27kb. When aligned to the hg38 reference genome^22^, the RNA002 dataset had 9,416,942 aligned reads (96.8% of total reads) spanning 12,932 annotated Gencode genes with at least 10 aligned reads, while the RNA004 dataset had 14,140,630 aligned reads (97.6% of total reads) spanning 13,680 annotated Gencode genes with the same filtering criteria (**see methods**).

We compared the alignment identity to the hg38 reference genome between RNA002 and RNA004. RNA002 data basecalled with Guppy v6.38 achieved a median identity of 90.65% for reads with at least 200 aligned bases, while RNA004 data basecalled with Dorado v0.8.1 achieved a median alignment identity of 98.67% for the same filtering criteria. When compared to older generations of basecalling software, we documented a continued improvement in alignment identity over the last 5 years (**Figure 1A**). When we computed the per-base substitution matrix, we noted that a reference base of C miscalled as U was the most prevalent miscall in both RNA002 and RNA004, with C-to-G and G-to-C as the least prevalent (**Figure 1B-C**). The rate of these global miscalls decreased by a factor of approximately 3 when transitioning from RNA002 to RNA004. Our data suggested that the highest per-base-substitution rate of 1.68% for C to U in RNA002 had decreased to 0.5% in RNA004. To assess the impact of read length on alignment identity, we binned reads into 500-nucleotide buckets starting at 0-500 (**Figure 1D**). We documented a higher variance in RNA002 alignment identity with a relatively steady median in each bucket, indicating that length had no discernible impact on alignment identity in light of the high variance. RNA004 showed a similar variance, with a strong skew (**Supplementary Table 2**) towards higher alignment identity for every bucket, but did observe a slight decrease in median alignment identity as lengths increased (**Supplementary Figure 2**). To better understand the overall increase in accuracy for RNA004, we computed the proportion of insertions and deletions (INDELS) in the aligned sequences. RNA002 had a median proportion of 7.19% INDELS, while RNA004 had a median proportion of 0.88% INDELS (**Figure 1E**). The reduction of INDELS in conjunction with a 3-fold improvement in per-base substitution rates helped to explain the increased alignment identity for RNA004.

**Figure 1.**
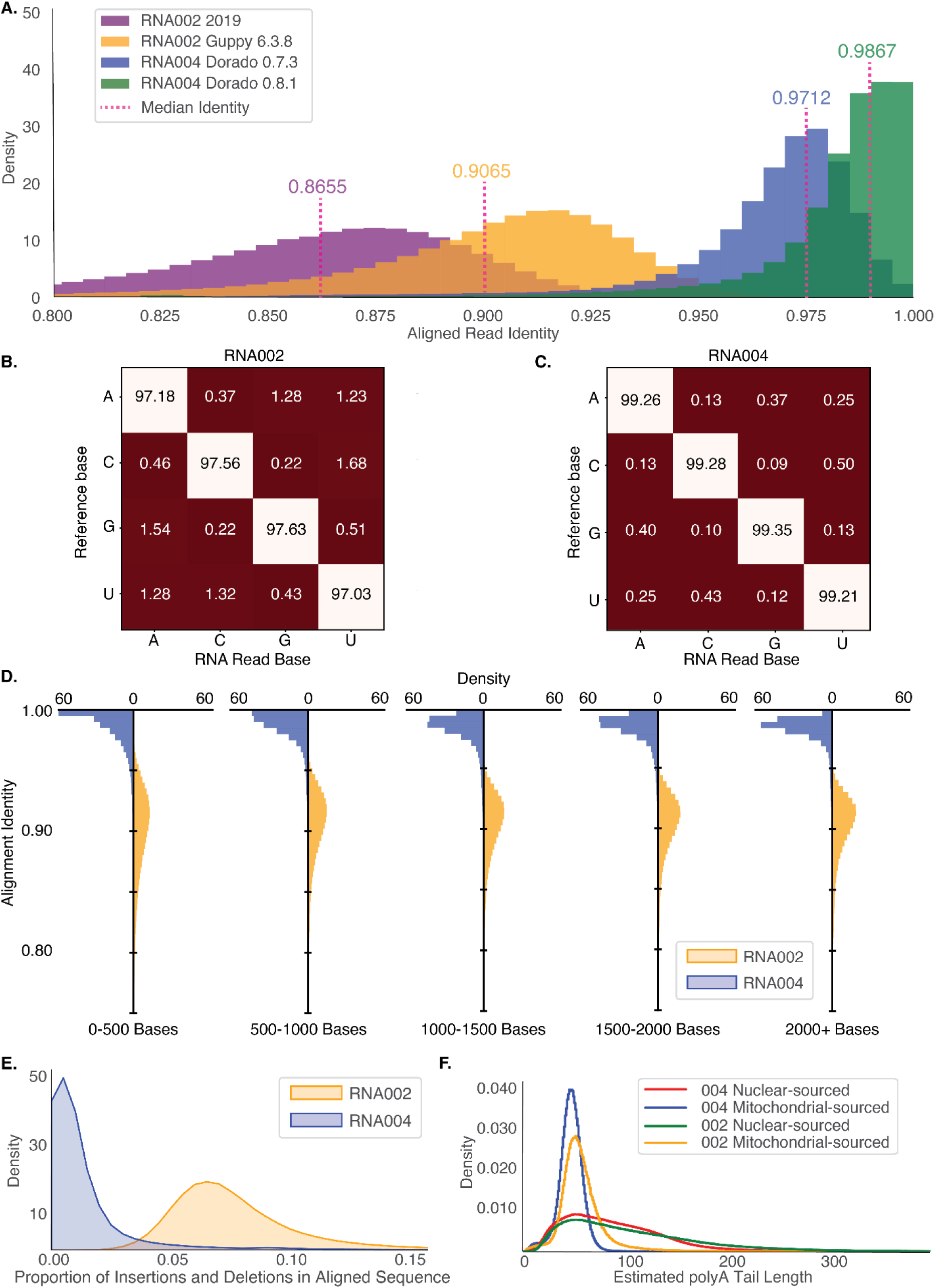
Comparison of RNA002 and RNA004 direct RNA sequencing performance. **A.** Aligned read identity was calculated as a percentage of total matches over total matches, mismatches, insertions, and deletions in the aligned region for RNA002 and RNA004. This calculation was repeated for multiple sequencing chemistries across multiple versions of basecallers all for the same extraction of GM12878 RNA. **B.** RNA002 base-specific substitution matrix calculating the rate at which, given a reference base, each possible canonical nucleotide was observed in aligned regions of reads. **C.** RNA004 base-specific substitution matrix using the same method as the RNA002 base-substitution matrix. **D.** Read identity histograms for RNA002 and RNA004 binned by aligned read length in 500 aligned nucleotide buckets. Aligned read length was calculated as the genomic distance between the first aligned base of the read and last aligned base of the read. **E.** Insertion and deletion proportion for RNA002 and RNA004 calculated as the number of insertions over total aligned bases and deletions over total aligned bases respectively. **F.** poly(A) tail estimates for mitochondrial and cytosolic transcripts for RNA002 (nanopolish) and RNA004 (Dorado).

Additionally, we investigated the RNA poly(A) tail length estimates between Nanopolish for RNA002 data and Dorado recently implemented native poly(A) tail estimation for RNA004 data. We grouped aligned reads into mitochondrial-sourced transcripts and genomic transcripts, with nuclear-sourced transcripts having an estimated poly(A) tail length median of 92.05 and 80 for RNA002 and RNA004, respectively. Mitochondrial-sourced transcript poly(A) length estimates had a median of 54.4 and 47 for RNA002 and RNA004, respectively. When we compared nuclear and mitochondrial-sourced transcripts between the two chemistries, we observed significantly different distributions for both populations (Nuclear-sourced transcript poly(A) Wilcoxon rank-sum, p < 2.22510^−308^, Mitochondrial-sourced transcript poly(A) Wilcoxon rank-sum, p < 2.22510^−308^) (**Figure 1F**). The difference in medians (RNA002 - RNA004) was 12.05 and 7.4 (13.091% and 12.963% reduction) for nuclear-sourced transcripts and mitochondrial-sourced transcripts, respectively. While there is a significant difference between the distributions of poly(A) tail lengths estimated by both chemistries, it is unclear which is more accurate. The preservation of the shape of the distributions between sequencing chemistries for two separate transcript types suggests that differential poly(A) tail analysis techniques employed in RNA002 experiments will yield similar results using the RNA004 sequencing chemistry.

### Number of Ψ sites substantially varies between RNA002 and RNA004 DRS data

Dorado additionally introduced a set of 4 RNA modification calling models. In this section, we compare two of these models (Ψ and m^6^A) against the RNA002 modification calling strategies for the same modifications.

As U-to-C mismatch was previously used as an indicator of uridine modifications (e.g., pseudouridine^5,6,23^), we compared U-to-C mismatch percentages for RNA002 and RNA004 for overlapping positions between the two datasets (**Figure 2A**). This analysis considered positions with at least 20 reads coverage and involved extracting positions where the reference base was U and the observed base was C. We computed the fraction of C reads over total reads and filtered for positions with at least one U-to-C mismatch and no genomic variants (**see methods**). The same method was used for calculating RNA004 U-to-C mismatch percentage. For 161,796 sites, RNA002 was 5+% higher in U-to-C mismatch. For 5,216 sites, RNA004 was 5+% higher in U-to-C mismatch. For 2,363,589 sites, the two chemistries were within 5% of each other (**Supplementary Figure 3**). To quantify differences, we calculated the Root Mean Square Error (RMSE), which measures the average deviation between two datasets. A lower RMSE indicates closer agreement between datasets, while a higher RMSE reflects greater differences. The RMSE between RNA004 and RNA002 U-to-C mismatch percentages was 0.0410.

**Figure 2.**
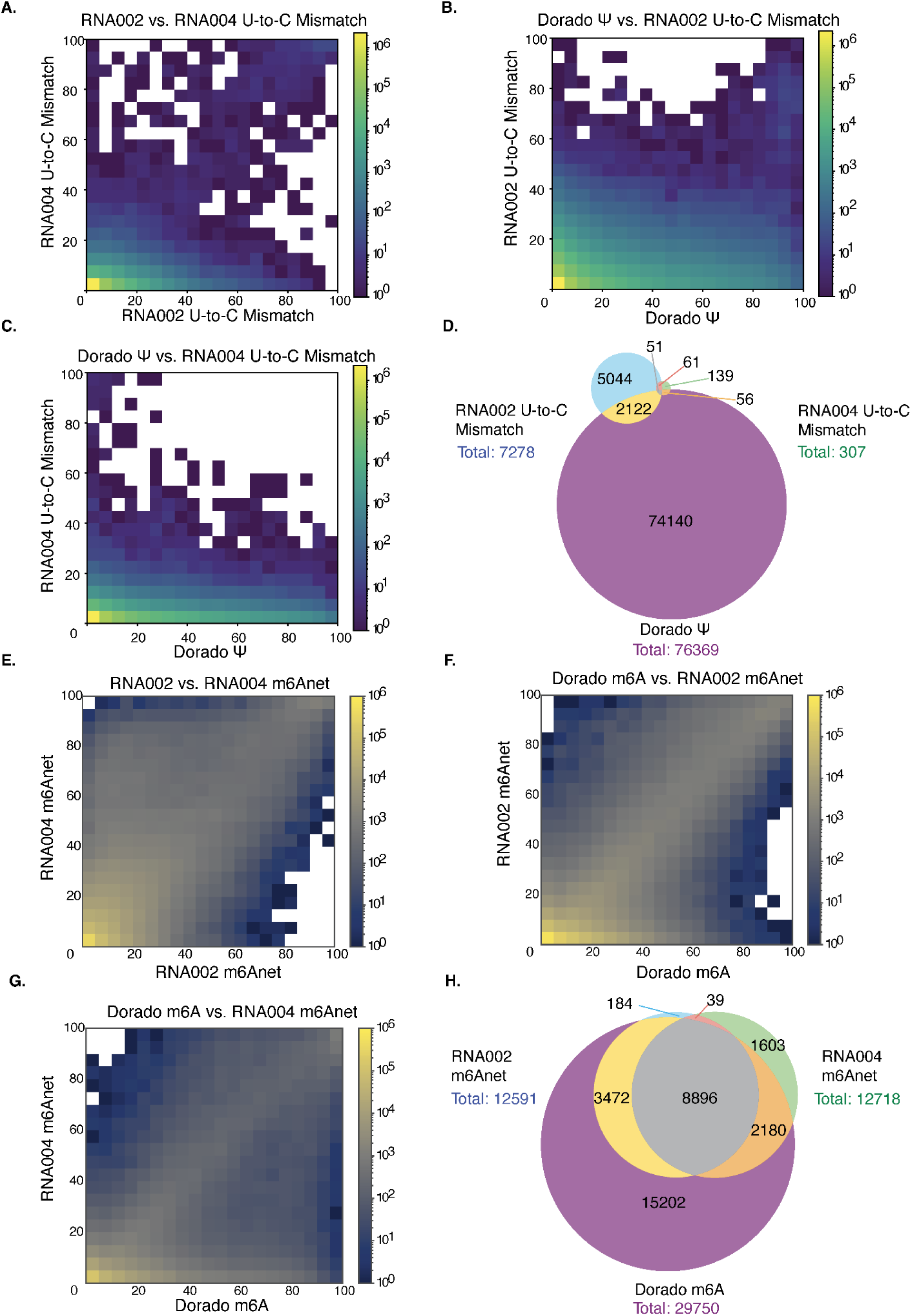
Comparison of Dorado Ψ and m^6^A models with the RNA002 modification calling strategies. **A.** Comparison of U-to-C mismatch percentage between RNA002 (x-axis) and RNA004 (y-axis) for reads with valid coverage of 20. **B.** Comparison of RNA002 U-to-C mismatch percentage (y-axis) and Dorado Ψ reported occupancy (x-axis) for reads with valid coverage of 20. **C.** Comparison of RNA004 U-to-C mismatch percentage (y-axis) to Dorado Ψ reported occupancy (x-axis) for reads with valid coverage of 20. **D.** Venn diagram showing the shared and unique sites between RNA002 U-to-C, RNA004 U-to-C and Dorado Ψ with at least 20% reported modification occupancy (or 20% U-to-C mismatch percentage) after intersecting all sites for valid coverage of 20 reads. **E.** Comparison of RNA002 and RNA004 m6anet reported occupancy. **F.** Comparison of RNA002 m6anet to Dorado m^6^A. **G.** Comparison of RNA004 m6anet to Dorado m^6^A. **H.** Shared and unique sites between RNA002 m6anet, RNA004 m6anet, and Dorado m^6^A.

We compared the U-to-C mismatch percentages for RNA002 to RNA004 Dorado Ψ model by considering overlapping positions between the two datasets with at least 20 reads coverage (**Figure 2B**). We also compared the distribution of RNA004 U-to-C mismatch percentage to RNA004 Dorado Ψ modification predictions (**Figure 2C**). In both instances, the density of sites was dispersed away from the line y=x, suggesting discordance between U-to-C mismatch and the Dorado model. The RMSE between Dorado Ψ modification and RNA002 U-to-C mismatch was 0.0887, and the RMSE between Dorado Ψ modification and RNA004 U-to-C mismatch was 0.1048.

For the intersection of valid sites in all three datasets, we used a Venn diagram to visualize the overlap of putative modification positions between datasets: RNA002 U-to-C mismatches, RNA004 U-to-C mismatches, and Dorado Ψ modification positions (**Figure 2D**). We limited the analysis to sites that had valid coverage (n=20) in all three datasets to control for batch variability. This decreased the number of sites considered (**Supplementary Figure 4A**). All three datasets were filtered to include positions with at least 20 reads and a 20% U-to-C mismatch or Dorado Ψ modification percentage. The genomic variants were excluded from the pool of sites. This comparison provides insights into the disagreement and agreement between different approaches. The U-to-C mismatch decreased from 7,278 positions in RNA002 to 307 sites in RNA004. This is while 76,369 of the eligible sites were detected by Dorado with at least 20% Ψ modified. Only 51 sites were shared among the three datasets. The decrease in the number of sites between RNA002 and RNA004 U-to-C mismatch indicates a sharp decrease in the miscall signal that previous studies have relied on. Compounding this with the lack of overlap between the Dorado Ψ caller and the RNA002 sites indicates that U-to-C mismatch is not a strong identification strategy using RNA004.

### Number of m^6^A sites shows higher concordance between RNA002 and RNA004 DRS data

We called m^6^A sites using m6anet for both RNA002 and RNA004 datasets. We converted the results to genomic positions and filtered for shared sites between both datasets (**Figure 2E**). We also compared Dorado m^6^A calls with the m6anet RNA002 dataset (**Figure 2F**) and m6anet RNA004 dataset (**Figure 2G**).

We used a Venn diagram to visualize the overlap of sites between datasets: RNA002 m6anet, RNA004 m6anet, and Dorado m^6^A modification positions (**Figure 2H**). As with Ψ, we limited the analysis to sites that had valid coverage (n=20) in all three datasets to control for batch variability. This again significantly decreased the number of sites considered, but provided a more robust foundation for comparison (**Supplementary Figure 4B**). All three datasets were filtered to include positions with at least 20 reads and 20% modification percentage. The sights for m6anet were further filtered for 90% probability of modification. This comparison provides insights into the disagreement and agreement between different approaches. The number of sites called by m6anet increased slightly from 12,591 positions in RNA002 to 12,718 positions in RNA004. In contrast, 29,750 m6A positions were called by Dorado. 8,896 positions were shared across the three datasets. The majority of m6anet sites are subset in the Dorado modification calls. This suggests that m6anet likely produced a more restrained set of modification calls, consistent with the algorithm’s dependence on pre-processing of reads through a method called resquiggling. While a purely signal-based approach such as Dorado will generally produce a larger range of callable sites, m6anet, for both 002 and 004, produced a high level of concordance with Dorado, indicating well trained models.

### Adjusting false positive rates for modification calls using IVT-derived canonical RNA DRS data

To improve the specificity of modification detection, we adjusted RNA004 Dorado modification calls using our whole genome IVT dataset^21^. Briefly, IVT RNA, composed only of canonical nucleotides, provides a biologically representative sequence context without modifications, making it an effective negative control. We used our pre-computed 9-mer false positive rates^21^ to adjust the Dorado modification calls by subtracting the 9-mer specific false positive rate from the biological and compared them against the original calls (**Figure 3A**). This strategy is broadly applicable for reducing the number of false positive calls and can be extended to future modification calling models for modifications described here as well as novel modification callers as they are developed.

**Figure 3.**
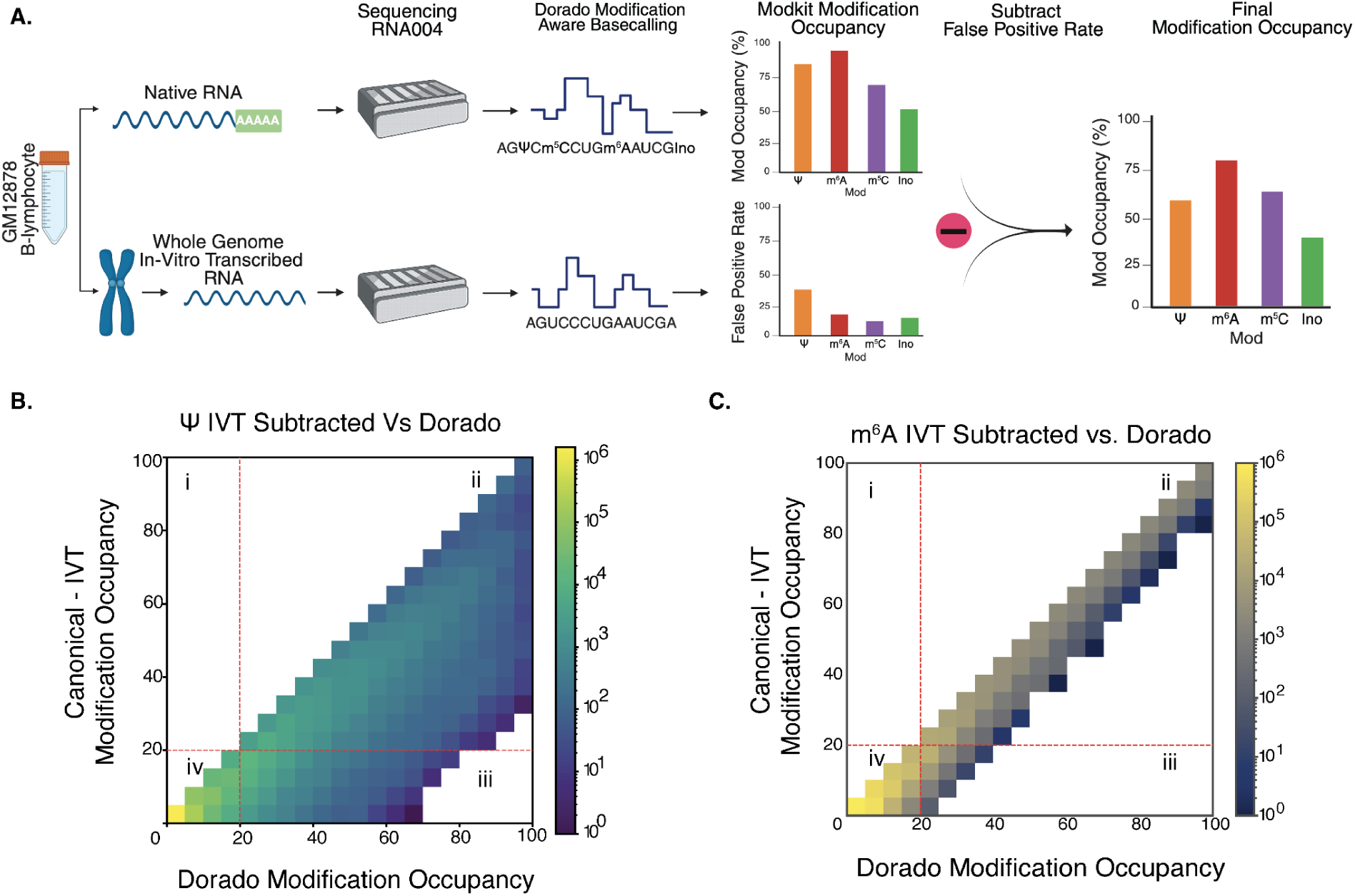
Adjusting false positive rates for modification calls using genomic DNA-based, IVT-derived canonical RNA. **A.** Schematic overview of the steps involved in comparing and subtracting IVT-derived false positive rates from native RNA modification calls obtained by Dorado. We used our pre-computed 9-mer false positive rates^21^ to adjust the Dorado modification calls by subtracting the 9-mer specific false positive rate from the biological and compared them against the original calls. **B.** Comparison of Dorado Ψ reported modification occupancy before subtracting IVT false positive rate (x-axis) and reported modification occupancy after subtracting IVT (y-axis). The red line indicates the 20% threshold and separates the 4 regions of the 2D histogram. **C.** Comparison of Dorado m^6^A reported modification occupancy before subtracting IVT false positive rate (x-axis) and reported modification occupancy after subtracting IVT (y-axis).

Each position was matched to its corresponding reference 9-mer, and IVT modification false-positive rate was looked up from the pre-calculated table. The IVT reported modification occupancy was then subtracted from the canonical raw Dorado reported modification occupancy to create an adjusted estimate of modification occupancy for each position. For biological Ψ sites, we compared the primary Dorado occupancy with the adjusted occupancy after subtracting the IVT-derived false positive rates. This approach aimed to reduce false-positive calls. Initially, 134,545 sites were detected by Dorado with at least 20 reads and a reported Ψ occupancy of 20%. After applying the IVT adjustment, this number was reduced to 74,043 sites (**Figure 3B and Table 4**). In the 2D histogram, we separated the sites into four regions based on the 20% threshold used throughout our analysis. Region “i” represents the sites (n=0) with < 20% original reported modification occupancy and ≥ 20% modification occupancy after subtracting IVT false positive rates, since the false-positive rate is >= 0, no sites gained estimated modification occupancy. Region “ii” contains the sites (n=74,043) with ≥ 20% reported modification occupancy both before and after IVT subtraction. Region “iii” includes the sites (n=56,577) with ≥ 20% original reported modification occupancy and < 20% occupancy after subtracting IVT. Region “iv” corresponds to the densest region of the histogram, containing 2,096,611 sites with < 20% reported modification occupancy both before and after IVT subtraction.

Similarly, the IVT occupancy was used for calling biological m^6^A sites by comparing the primary Dorado reported modification occupancy with the adjusted occupancy after subtracting the IVT-derived false positive rates. The dataset originally contained 185,714 sites where Dorado had 20 reads and called m^6^A with ≥ 20% occupancy. After utilizing the IVT correction, this number was reduced to 138,670 (**Figure 3C and Table 4**). As was with Ψ, the figure is separated into four regions based on the 20% threshold used throughout our analysis. Dorado m^6^A showed a similar pattern, with region “i” (n=0), region “ii” (n=138,670), region “iii” (n=47,044), and region “iv” (n = 5,140,533). The IVT adjustment for putative m^6^A sites increased overlap with GLORI-Seq (**Supplementary Table 3**).

### Orthogonal validation of select Dorado modification calls using synthetic oligonucleotide DRS data

To further evaluate the performance of Dorado modification calling for detecting Ψ and m^6^A on known modification states, we designed 6 synthetic oligonucleotides and their corresponding *in vitro* transcribed pair with a modification at a known position. 3 of these oligos were designed for Ψ and 3 for m^6^A (**Table 1 & 2, see methods**). For each of the synthetically modified oligonucleotides, a true negative oligonucleotide was generated using IVT (**see methods**).

**Table 1.** Modification heuristics for RNA002 and RNA004 for selected positions with Ψ predictions.

| Name | U-to-C Mismatch (RNA002) | Dorado occupancy (RNA004) |
| --- | --- | --- |
| $\Psi$_1 | 100% | 100% |
| $\Psi$_2 | 100% | 0% |
| $\Psi$_3 | 0% | 100% |

**Table 2.** Modification heuristics for RNA002 and RNA004 for selected positions with m^6^A predictions.

| Name | Dorado occupancy - RNA002 m6anet occupancy | Delta Percentile Range |
| --- | --- | --- |
| $m^6A$_1 | -20.91% | 0 - 5th |
| $m^6A$_2 | 23.47% | 95 - 100th |
| $m^6A$_3 | 10% (Both $\geq 90\%$) | 65 - 70th |

The libraries were prepared using the DRS protocol and the latest SQK-RNA004 kit for the 12 constructs (synthetic modification-bearing oligonucleotides and canonical IVT-derived RNA). The raw data were basecalled using Dorado v0.8.1 with Ψ and m^6^A modification calling models, yielding 56,443,656 reads. Filtering for primary alignment resulted in 33,141,855 reads (57% of all aligned reads). We further filtered for full length and identification of a barcode sequence to ensure their identity, which resulted in 15,745,700 reads (**see methods**). The number of full-length reads for each oligonucleotides is detailed in **Table 3**. ONT’s Modkit tool was run on the filtered aligned BAM file with the modification tags. We compared the modification occupancy of our known modified sites with their IVT pair. We also evaluated the performance of traditional modification detection strategies on these data.

**Table 3.** Number of full-length reads for all oligonucleotides and Dorado performance for each pair.

| Oligomer Name | Mod Reads | IVT Reads | Balanced Accuracy | Accuracy | Precision | Recall | F1 Score |
| --- | --- | --- | --- | --- | --- | --- | --- |
| $\Psi_1$ | 2,236,489 | 362,850 | 0.9839 | 0.9806 | 0.9972 | 0.9782 | 0.9876 |
| $\Psi_2$ | 1,851,854 | 495,933 | 0.9710 | 0.9638 | 0.9950 | 0.9574 | 0.9759 |
| $\Psi_3$ | 1,340,557 | 947,062 | 0.9650 | 0.9644 | 0.9859 | 0.9441 | 0.9646 |
| m6A_1 | 2,071,393 | 159,672 | 0.9851 | 0.9868 | 0.9989 | 0.9869 | 0.9929 |
| m6A_2 | 1,804,781 | 756,602 | 0.9616 | 0.9486 | 0.9965 | 0.9316 | 0.9629 |
| m6A_3 | 1,792,189 | 1,080,786 | 0.9737 | 0.9726 | 0.9867 | 0.9693 | 0.9779 |

For the three known Ψ positions, Dorado occupancy ranged from 94% to 97%, while the U-to-C mismatch percentage varied between 2% and 22% at the same sites (**Supplementary Figure 5A**). In the IVT pairs, Dorado occupancy remained around 1% and the U-to-C mismatch percentage was approximately 0%. We compared Dorado m^6^A calls with m6anet for these synthetic oligonucleotides (**Supplementary Figure 5B**). Notably, 99.83% of reads were disqualified from analysis by m6anet due to upstream filtering by eventalign. Consequently, only one of the three known synthetic oligonucleotides (m^6^A_1) was identified as modified with greater than 90% probability according to m6anet’s modification criteria.

Additionally, we calculated performance metrics for Dorado performance on the synthetic oligonucleotides (**Table 3**). The Dorado Ψ model achieved both accuracy and F1-score in the range of 96% to 98%, while the Dorado m^6^A model showed accuracy between 94% and 98% and an F1-score ranging from 96% to 99%.

We analyzed the modification caller’s confidence at a per-read per-site basis for each of the oligonucleotides. We compared the modification-bearing oligonucleotide with the IVT oligonucleotide for both Dorado Ψ and m^6^A, at the known modified position and two reference bases surrounding that position (**Figure 4. A-C & D-F**).

**Figure 4.**
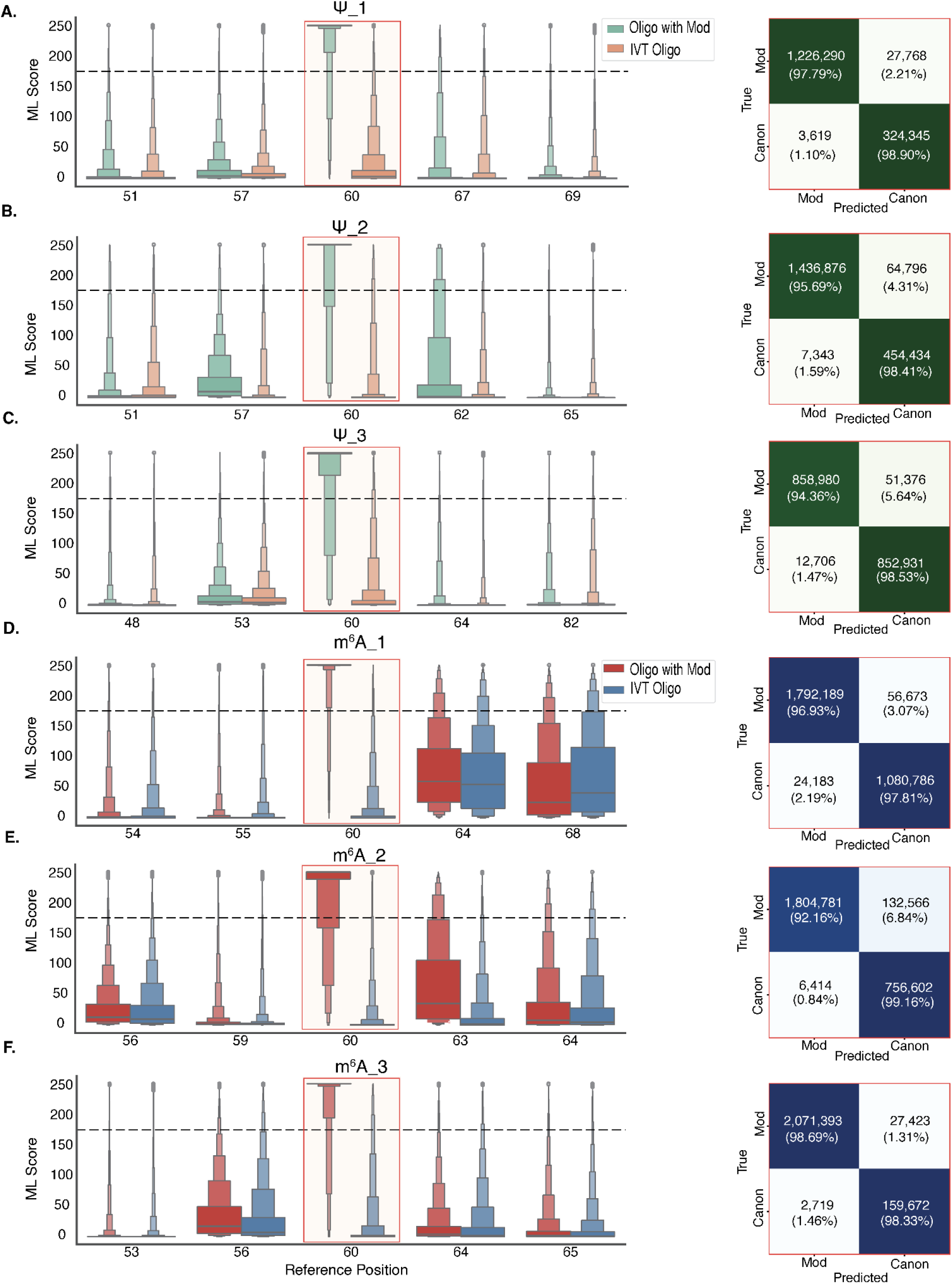
Evaluation of Dorado modification calling performance on known synthetic modification-bearing and conconical IVT oligonucleotides. **A.** Boxen plot comparing the raw Dorado modification likelihood (ML) scores at the known central modified site (60) and 2 reference U sites surrounding it for Ψ_1, with confusion matrix of prediction at the modified site in Ψ_1. **B.** Boxen plot comparing the raw ML tag scores at the known central modified site (60) and 2 reference U sites surrounding it for Ψ_2, with confusion matrix of prediction at the modified site in Ψ_2. **C.** Boxen plot comparing the raw ML tag scores at the known central modified site (60) and 2 reference U sites surrounding it for Ψ_3, with confusion matrix of prediction at the modified site in Ψ_3. **D.** Boxen plot comparing the raw ML tag scores at the known central modified site (60) and 2 reference A sites surrounding it for m^6^A_1, with a confusion matrix of prediction at the modified site in m^6^A_1. **E.** Boxen plot comparing the raw ML tag scores at the known central modified site (60) and 2 reference A sites surrounding it for m^6^A_2, with a confusion matrix of prediction at the modified site in m^6^A_2. **F.** Boxen plot comparing the raw ML tag scores at the known central modified site (60) and 2 reference A sites surrounding it for m^6^A_3, with a confusion matrix of prediction at the modified site in m^6^A_3.

We evaluated Dorado performance on these known modified conditions and their IVT pairs (**Figure 4. A-C & D-F**). True positives are where the modified base was correctly identified as modified (left upper box of the confusion matrix). False negatives are where the modified base was incorrectly identified as canonical (right upper box of the confusion matrix). False positives are where the canonical base was incorrectly identified as modified (left lower box of the confusion matrix). True negatives are where the canonical base was correctly identified as canonical (right lower box of the confusion matrix). Dorado identified 94% to 98% of true positives, with 1% to 2% false positives across the six oligonucleotide pairs.

### Dorado calling for Ψ, m^6^A, m^5^C, and Inosine

In conjunction with Ψ and m^6^A, we used Dorado to call m^5^C and inosine. Across all four modifications, Dorado called 419,757 putative sites. We identified 285,317 high-confidence sites that each had 20 aligned reads with a modification occupancy of 20% or higher when adjusted for the 9-mer specific false positive rates (**Table 4; see Methods**). All four modifications showed a generally decreasing number of sites as occupancy cutoffs increased, with m^6^A showing similar numbers of sites for 60-70%, 70-80%, 80-90% and 90%+ cutoffs. (**Figure 5A**). We note that m^6^A had 5,818 sites with a reported occupancy of 90-100%. In contrast, Inosine had 36 sites for the same occupancy threshold. To better understand the distribution of modifications at a more granular level, we collated modification sites with at least 20 reads and 20% occupancy for each gene as defined by gencode v43. We documented 12,116 genes with more than 1 instance of at least one modification type (**Figure 5B**), with some genes harboring more.

**Figure 5.**
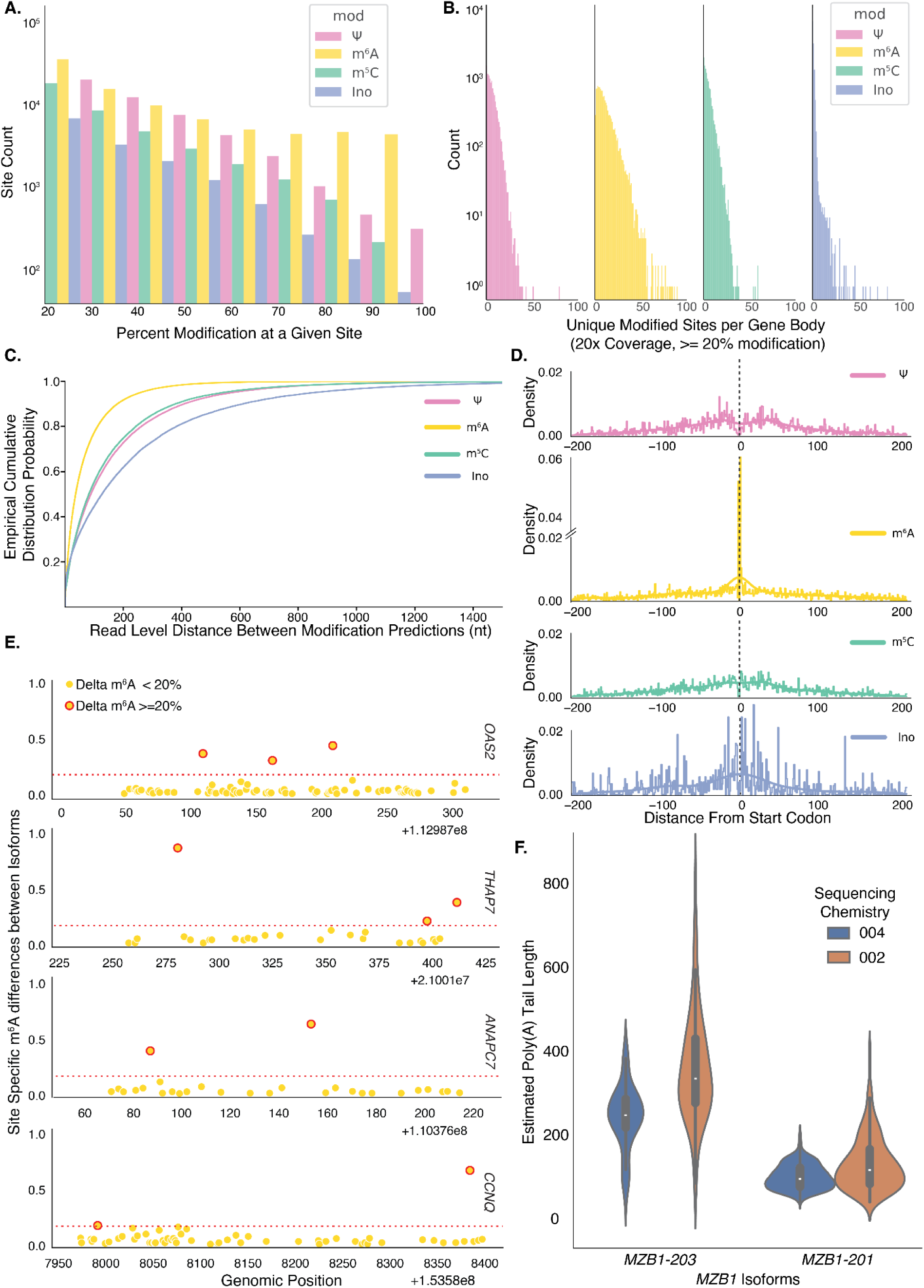
Dorado modification calling for Ψ, m^6^A, m^5^C and inosine. **A.** Count of sites with 20 or more reads, 20% modification occupancy after subtracting 9-mer specific false positive rates. Counts per modification type were binned into 10% estimated modification occupancy windows starting at 20-30% occupancy and increasing by 10% increments. **B.** Counts of genes binned by the number of occurrences of the given modification across the entire gene body. Modifications were only counted if they met the same 20 read coverage with 20% corrected modification occupancy. **C.** Empirical cumulative distribution of distance between RNA modifications on a given read. Each line shows the cumulative distribution function for a single RNA modificaiton type, and the distances between modifications of that same type. **D.** Density of modifications in proximity to the gencodev43 start codon positions. Densities were calculated exclusively for the +/− 200nt window around the start codon, and the start codon itself was excluded from analysis. **E.** Selection of four exemplar genes with high levels of isoform specific m6A expression. Each dot on the figure represents a common site between at least two isoforms of the given gene. The x-axis denotes genomic position, while the y-axis represents the maximal m^6^A percent occupancy distance between any two isoforms of that gene at that position. Positions that had a delta of greater than or equal to 20 are circled in red. **F.** An exemplar gene with significant Isoform-specific differences in poly(A) tail estimation for RNA002 and RNA004. Estimates for RNA002 were produced with nanopolish polya, while RNA004 estimates were produced with RNA004.

**Table 4.** Number of Dorado modification sites before and after subtraction of IVT false positive rates.

| Mod | Number of sites <u>before</u> subtracting IVT ($\geq 0\%$, $\geq 20\times$ coverage) | Number of sites <u>before</u> subtracting IVT ($\geq 20\%$, $\geq 20\times$ coverage) | Number of sites <u>after</u> subtracting IVT ($\geq 20\%$, $\geq 20\times$ coverage) |
| --- | --- | --- | --- |
| $\Psi$ | 3,625,185 | 134,649 | 74,043 |
| $m^6A$ | 5,326,257 | 185,714 | 138,670 |
| $m^5C$ | 3,183,370 | 75,087 | 57,285 |
| Inosine | 2,830,935 | 24,307 | 18,615 |

In addition to gene-level differences, we calculated the distance between modifications of the same type at the read level; m^6^A had a higher probability of occurring in close proximity to other m^6^A modifications on a read when compared to the distance between; Ψ and Ψ, m^5^C and m^5^C, inosine and inosine (**Figure 5C**). Given the higher abundance of m^6^A calls this was not an unexpected result. We catalogued the site-specific distribution of modifications across gene bodies. Prior work has demonstrated that the localization of modifications around specific gene components can impact translation^24–27^. We investigated broad trends for the deposition of these four modifications around the start codon (**Figure 5D**). Only m^6^A showed a clear pattern of hypermodification in proximity to the start codon. This relative abundance of modifications occurred on both the 5′-UTR and the coding sequence side of the start codon. We identified isoform-specific patterns of modifications using Isoquant^28^ and Modkit. We documented isoform-specific occupancy differences as high as 80% for m^6^A (*OAS2*), with isoform pairs of other genes ranging in the 25-75% range (**Figure 5E**). We verified that Dorado poly(A) estimates were concordant with the isoform-specific poly(A) tail length differences that we and others had previously shown. To that end, we computed isoform-specific polyA tail estimations for the *MBZ1* gene transcripts. For the RNA004 *MZB1* transcripts, we were able to identify a significant difference in the poly(A) tail length distributions (Kolmogorov–Smirnov, p=4.24×10^−78^) for the isoforms *MZB1-203* and *MZB1-201*. We were able to duplicate this finding using RNA002 (p=9.02×10^−43^), indicating that poly(A) tail length estimation remained informative across chemistry updates (**Figure 5F**).

### RNA004 DRS data predict co-occurrence of multiple RNA modifications

To better understand how multiple types of modifications occur together, we calculated the co-observation rate between pairs of modifications. Here, we defined the co-observation rate as the proportion of genes with a given modification B, given modification A on the gene body. This co-observation rate was calculated pairwise for the four modifications. The co-observation rate for Inosine, the least frequent modification of our set, given any of the other three modifications was at least 39.4%. Between Ψ, m^5^C, and m^6^A, the lowest co-observation rate was higher at 79.6%. Additionally, for genes with at least one m^6^A call, we documented a 100% co-observation rate for a Ψ call in this dataset (**Figure 6A**). Given that Ψ and m^6^A both had putative sites in excess of the number of observed genes, it was not surprising that if an m^6^A was observed, so was a Ψ. Due to the high level of co-observation of modifications across gene bodies, we wanted to understand the relationship between read length and the number of modifications called by Dorado. The relationship between read length and the number of modifications called across the entire read produced two distinct populations, each with an independent but clear linear trend (**Figure 6B**).

**Figure 6.**
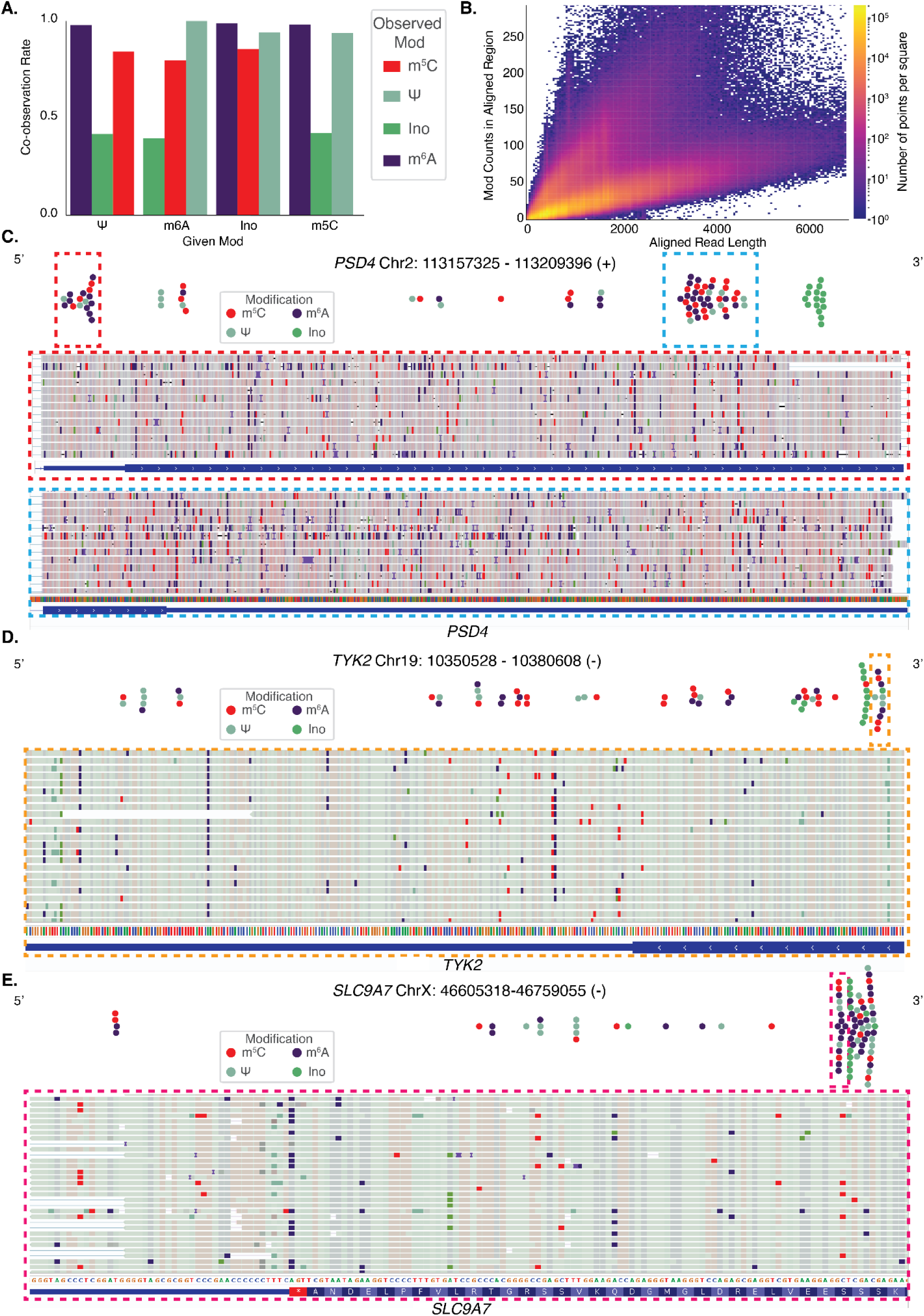
Co-occurrence of multiple RNA modifications. **A.** Co-observation rates, calculated as the probability of observing modification X given modification Y, across the entire set of sequenced gene bodies. All modification calls that occurred within 5nt of each other were excluded from this analysis to limit the impact of modification associated miscalls affecting the analysis. **B.** Relationship between aligned read length and the number of predicted mods at a per-read level. Modification counts per read were calculated using the estimated threshold from the modkit pileup analysis. **C-E.** Exemplar genes with putative modification sites (>= 20 reads, >= 20% IVT adjusted modification occupancy) for each of the four target modifications, with supporting IGV screenshots.

This behavior has several possible explanations including reads with abnormal ionic current signal, that may have produced a higher number of modification calls. Or it is possible that some transcripts are genuinely modified to a higher degree than their counterparts. These observations warrant further investigation and validation, and will likely yield biological or computational insight for the community.

Because inosine had the least number of Dorado calls, we decided to investigate why m^6^A and Inosine, as a pairwise set of modifications, occurred at high quantities over a handful of genes. When we investigated the sites individually, we found that these regions were low complexity A/G rich regions. We investigated the mapping quality and Phred scores of reads in the low complexity sequence occurring in an intronic region of *KIRREL3* (chr11:126,923,084-126,923,447) and found that the Phred scores, modification scores and mapping qualities were systematically low in comparison to randomly selected positions from the transcriptome (**Supplementary Figure 6**). We hypothesize that these reads were alignment artifacts originating from poor-quality reads. The modification caller’s confidence in these regions produced a pattern in Modkit where most of the reads, modified or canonical, fell below the fail threshold. With reads falling below the fail threshold being withheld from the coverage and percent modification calculations, Modkit reported a high modification status on a subset of the low-quality data. These alignment artifacts showed abnormally high quantities of Inosine and m^6^A but only accounted for a small number of genes with multiple modification types deposited in unison.

With adjustment for false positives, we examined other genes with high rates of multiple types of modifications to determine if they were also the product of low complexity regions, poor alignments, or another potential source of error. However, our analysis identified a variety of genes with multiple modifications of various types deposited across gene bodies (**Figure 6C-E)**. Some of these genes had as many as 139 reported modification sites.

## Discussion

In 2024, ONT released its new RNA004 chemistry, which claimed to offer higher yield and improved accuracy compared to the older RNA002 chemistry. Alongside this update, ONT also enhanced its basecaller Dorado, which now supports *de novo* detection of four RNA modifications^15^: Ψ, m^6^A, m^5^C, and Inosine. Dorado enabled site-level modification analysis without requiring negative controls.

We analyzed RNA002 and RNA004 DRS data for poly(A) selected RNA from GM12878 B-lymphocyte cell line. We expanded our analysis to compare all four RNA modifications that were detected by Dorado. We compared orthogonal modification identification methods for Ψ (U-to-C mismatches) and m^6^A (m6anet). For both datasets, we observed site-specific differences in U-to-C mismatch and m^6^A occupancy despite originating from the same RNA sample. Traditionally, U-to-C mismatches were used as a method for detecting Ψ, but we observed significant discrepancies between U-to-C mismatch ratios and the Dorado Ψ modification caller.

We also designed six synthetic oligonucleotides with known modifications (three for Ψ, three for m^6^A) alongside a canonical IVT pair. We evaluated Dorado modification calling performance on the known modification condition. Dorado was able to identify the modified Ψ and m^6^A with an accuracy of 0.94-0.98% and an F1 score of between 0.96 to 0.99, with a false positive rate of 1-2%. This is while the U-to-C mismatch percentage for the known Ψ sites varied between 2% and 22%. While Dorado outperformed other modification detection strategies, it exhibited some false positive calls, highlighting the need for further validation and refinement. These refinements may involve manually inspecting the modification calls (sequence context and region), considering a suitable coverage and occupancy threshold in Dorado result to reduce false positives. To increase specificity of Dorado modification calls, we adjusted modification occupancy by subtracting genomic IVT false positive rates.

In conjunction with DRS long read lengths that allow for analysis of allele-specific expression at gene and isoform levels^29,30^, this modification information provides a unique and powerful lens to study RNA biology. However, without cross-validation through orthogonal methods (LC-MS/MS, enzymatic knockdowns, etc.) these modification predictions are putative at best.

## Conclusion

Nanopore DRS RNA004 update has demonstrated improved median accuracy (>98.6% in this study) and the ability to detect four types of RNA modifications (Ψ, m^6^A, m^5^C and inosine). Comparing RNA modification detection methods between RNA002 and RNA004 DRS data revealed significant discrepancies. This suggests that methods originally developed for RNA002 DRS data may not directly translate to RNA004, especially U-to-C mismatch for the identification of pseudouridine. By incorporating false positive controls such as the genomic IVT RNA strategy we detailed, further confidence can be placed in the prediction of positional modifications. Additional caution should be exercised when investigating putative modifications that occur in low complexity regions or in close proximity to other putative modifications.

Our analyses further demonstrate that it is possible to detect multiple RNA modifications simultaneously, each at a per-read, per-site level using the RNA004 chemistry. However, we argue that validating these modifications using orthogonal techniques such as chemical assays, synthetic and biological controls, and LC-MS/MS increases the confidence in biological conclusions drawn using nanopore modification detection. With supporting orthogonal validation, Nanopore DRS is a powerful tool for studying RNA modifications and their interplay in biological mechanisms and function.

## Materials and Methods

### GM12878 cell culture and RNA isolation

The poly(A) RNA used in this study was sourced from a tube of poly(A) RNA that was previously used in a 2019 study^1^. GM12878 cell culture, RNA extraction, and poly(A) RNA enrichment details have been previously described^1^. Briefly, low passage (passage 4) cells were cultured, and total RNA was isolated using Trizol-Chloroform extraction. This was followed by poly(A) RNA enrichment using NEXTflex Poly(A) Beads (BIOO Scientific cat. no. NOVA-512980) and stored at −80°C. This poly(A) RNA was sequenced using RNA002 chemistry previously^1^. For this study, we rebasecalled the RNA002 dataset using Guppy v6.3.8 (hac model). We sequenced 100 ng of poly(A) RNA from the RNA preparation from the previous study using RNA004 chemistry under standard conditions. These data were basecalled using Dorado^31^ basecaller v0.8.1 with the super accuracy model (rna004_130bps_sup@v5.1.0). Basecalling was repeated 3 times, once for each set of modifications (Ψ, m^6^A/Inosine, and m^5^C).

~~~
dorado basecaller sup --modified-bases inosine_m6A
dorado basecaller sup --modified-bases m5C
dorado basecaller sup --modified-bases pseU
~~~

For both RNA002 and RNA004, reads were aligned to the human reference genome (hg38.p13) using minimap2^32^, then sorted and indexed using Samtools.

~~~
minimap2 -uf -k 14 -ax splice | samtools sort -o
samtools index
~~~

### Biological Modification Calling

#### RNA002 U-to-C mismatch

To calculate U-to-C mismatches, we first ran Pysamstats^33^ on RNA002 aligned, filtered, and sorted BAM file using the GRCh38 genome reference.

~~~
pysamstats --type=variation_strand
--fields=“chrom,pos,ref,deletions,deletions_fwd,deletions_rev,insertions,insertions_fwd,insertions_rev,A,A_fwd,A_rev,C,C_fwd,C_rev,T,T_fwd,T_rev,G,G_fwd,G_rev”
~~~

To identify U-to-C mismatches, we first filtered for valid positions that have at least 20 reads. We analyzed positions where the reference base was T (representing U in RNA) and the observed base in sequencing reads was C. This calculation was repeated for the reverse strand. U-to-C mismatch was calculated as the number of C bases observed over the total number of bases at these positions. We limited our analysis to positions where this fraction was greater than zero (indicating at least one U-to-C mismatch). We then calculated the percentage contribution of each base (C, G, A) at these positions.

To ensure that U-to-C mismatches were not confounded by genomic variants, we used Clair3^34^, a variant caller designed for long-read sequencing, to exclude genomic variants from all analyses. We ran Clair3 on sequenced and aligned genomic DNA from GM12878 with the r1041_e82_400bps_sup_v500 model to identify positions with potential genomic variations. All variant positions were then excluded from the U-to-C mismatch calculations, eliminating genomic variation as a possible source of error. Ensuring that only genuine mismatches were considered in our analysis.

### RNA004 Ψ modification calling and U-to-C mismatch

The same steps used for calculating U-to-C mismatches in the RNA002 dataset were applied to the RNA004 dataset. This included using Pysamstats, extracting positions where the reference base was T (U in RNA) and the observed base was C, computing the fraction of C reads filtering positions with at least one U-to-C mismatch, and calculating the percentage contribution of each base. This was repeated for the reverse strand. Additionally, genomic variants identified using Clair3 were excluded to ensure that only mismatches were considered.

For identifying putative Ψ sites in RNA004, the sorted and filtered BAM file originating from the Dorado Ψ modification caller was analyzed with ONT’s Modkit^35^ tool to extract predicted modification information.

~~~
modkit pileup --motif T 0
~~~

Only positions supported by at least 20 reads were considered in the final analysis of Modkit results.

### m^6^A Calling with m6anet and Dorado

To compare the RNA002 chemistry, we used m6anet^12^ to detect m^6^A in DRS data. Reads were aligned to the Gencode v43 transcript^36^ reference using Minimap2. The aligned BAM file was subsequently sorted, filtered to include only primary reads, and indexed using Samtools. Nanopolish^37^ eventalign was used to produce event-level alignments for use with m6anet.

~~~
nanopolish eventalign --scale-events --signal-index
~~~

We used m6anet to identify putative m^6^a sites in the RNA002 data. m6anet’s HCT116_RNA002 model was used for inference. We called m^6^A sites in the RNA004 data using m6anet as well. The same series of steps were followed to generate these calls, with the exception of f5c^38^ being used to produce the event-level alignments since nanopolish has not been updated for RNA004 data.

~~~
f5c eventalign --rna --signal-index --scale-events
~~~

m^6^a was called with Dorado’s ino_m6a model for the RNA004 data. Site-level information was tabulated using modkit pileup.

### Modification occupancy adjustment and re-filtering with Whole Genome IVT Data

Using a 9-mer false positive rate table^21^, we calculated the reported modification occupancy of each site in the modkit pileup. After subtracting the false positive rate we refiltered for sites that continued to have a reported modification occupancy over 20% and a minimum of 20 valid (Modified or Canonical) read coverage.

### Selection of Sites for Designing Synthetic Oligonucleutides

To evaluate the performance of Dorado modification calling and the traditional method used for detecting Ψ and m^6^A on known modification states, we designed 6 synthetic oligonucleotides and their corresponding *in vitro* transcribed pair.

### Ψ site selection criteria

For the RNA002 dataset, we matched positions and chromosomes between Pysamstats U-to-C mismatch data and Dorado Ψ predictions. We applied filters requiring at least 20 reads and categorized positions into three groups:

- **Ψ_1**: High U-to-C mismatch & High Dorado Ψ (both 100%)
- **Ψ_2**: High U-to-C mismatch & Low Dorado Ψ (U-to-C = 100%, Dorado Ψ = 0%)
- **Ψ_3**: Low U-to-C mismatch & High Dorado Ψ (U-to-C = 0%, Dorado Ψ = 100%)

We then analyzed the resulting sequences, excluding positions located in homopolymer regions. The identified regions were visually inspected using the Integrative Genomics Viewer (IGV)^39^, ensuring that no designed oligonucleotide spanned an exon junction. Further structural and thermodynamic analyses were performed, including secondary structure prediction using RNAfold and ΔG (Gibbs free energy) calculations to assess sequence stability, dimer formation, and homodimer probability analysis to minimize unwanted interactions, melting temperature, GC content, and other physicochemical properties analyzed using Integrated DNA Technologies (IDT) tools. The three sequences we selected met all these criteria.

### m^6^A site selection criteria

We calculated site-specific occupancy differences between Modkit’s output and the genomic positions calculated from m6anet’s output. This allowed us to select a set of sites that represented large differences in m^6^A occupancy as calculated by the two technologies:

- **m^6^A_1:** High m6anet & Low Dorado (0^th^ - 5^th^ percentile of delta occupancy)
- **m^6^A_2:** Low m6anet & High Dorado (85^th^ - 95^th^ percentile of delta occupancy)
- **m^6^A_3:** High m6anet & High Dorado (>= 90% modified for both)

#### Synthetic oligonucleotide design

Based on the selection criteria mentioned, we designed six 30-base-long sequences, three for Ψ and three for m^6^A. A known modification was placed in the middle of each sequence. To be able to sequence the selected synthetically modified oligonucleotides, we performed a splinted ligation strategy, attaching a 3′ and 5′ oligonucleotide (**Supplementary Figure 1**, **Supplementary Table 1**). The 3′ oligonucleotides included a 3′ terminal sequence (polyadenylated tail) that made the full structure compatible with ONT’s DRS sequencing kit. The 3′ (60 bases) and 5′ oligonucleotide (46 bases) extended the total sequence length to 136 bases. A fully complementary single-stranded DNA template (ssDNA) was reverse-transcribed to hybridize with all oligonucleotides.

### Synthetic oligonucleotide experimental protocol

During the annealing process, RNA oligonucleotide, ssDNA, 3′ oligonucleotide, and 5′ oligonucleotide (1 μL of 100 μM each) were mixed with 6 μL of annealing buffer (10 mM Tris-HCl, pH 8.0, 50 mM NaCl) to a final volume of 10 μL at 10 μM concentration per tube. The mixture was then incubated at 75°C for 2 minutes and reduced to 25°C at a rate of 2°C per minute before being held at 4°C to facilitate annealing. For the ligation step, T4 RNA Ligase 2 (NEB #M0239) was used to ligate the adapter. The reaction mixture consisted of 5 μL of 10 μM of each construct (from the annealing step), 4 μL of 5X Quick Ligase Buffer (NEB # B6058), 8 μL of ultrapure water, and 3 μL of T4 RNA Ligase 2, bringing the total volume to 20 μL. The mixture was incubated at room temperature for 1 hour. This was followed by a 1.5X SPRI beads (Beckman Coulter # B23318) clean up with 2 wash steps using 70% ethanol.

We utilized IVT to synthesize a true negative control to compare to the synthetic oligonucleotides. We designed 6 DNA oligonucleotides that contained the reverse complement sequence to the synthetic oligonucleotides in the 5′ and modified sections (**Supplementary Table 1**). To differentiate the true positive modified and true negative IVT from a mixed sample, we added a 15-nucleotide barcode near the 3′ end of the IVT template oligonucleotide. We also included a T7 polymerase targeting site at the 5′ end of the DNA reverse complement oligonucleotide. The complementary strand was generated using a one-cycle PCR reaction. For each template, the reaction mixture included 1 μL of 10 mM DNA template, 1 μL of 10 mM dT Primer, 12.5 2x Long AMP Taq (NEB #M0323), and 10.5 μL of RNase-free water, which were combined in a separate tube for each template. Using the HiScribe NEB T7 High Yield RNA synthesis kit (#E2040), we transcribed true negative controls for our modified oligonucleotides. After *in vitro* transcription, reactions were treated with DNase I (NEB #M03030) for 15 min. Concentrations of all RNA oligonucleotides were determined with NanoDrop.

Following the DRS library preparation protocol with SQK-RNA004 kit^40^, the libraries for 12 oligonucleotides were made in a single tube (a total of 9.5 uL RNA with 32 ng of each construct). The RNA RC, RNaseOUT, and the optional reverse transcription step were skipped and the protocol was followed through sequencing. The library was sequenced using an RNA004 flow cell on a PromethION platform for 72 hours. The resulting Pod5 files were basecalled with Dorado 0.8.1 with modification calling enabled for Ψ and m^6^A separately.

~~~
dorado basecaller sup,pseU --modified-bases-threshold 0 –-no-trim
dorado basecaller sup,inosine_m6A --modified-bases-threshold 0
--no-trim
~~~

We used the complete sequence of all 12 oligonucleotide to create the reference. This reference was utilized with BWA-MEM for alignment.

~~~
bwa mem -W 13 -k 6
~~~

The resulting Sam file was converted to Bam, sorted and indexed using Samtools. Then, we copied the modification tags (MM and ML) from the Dorado unaligned BAM to the bwa-mem aligned BAM. Utilizing the barcode attached to the end of the IVT oligonucleotides, we filtered the data to only include reads that aligned between the 5′ end of the synthetic 30-nucleotide sequence and the 3′-end of the barcode. This process was performed as well at positions that correspond to those indices in the synthetic modified oligonucleotides.

### Statistical analysis

Per read identity was calculated as the number of matches divided by the number of matches + mismatches + insertions + deletions for reads aligned to gencode.v43. We additionally limited identity calculations to reads with an aligned length of at least 200 (reference end - reference start).

To calculate the base substitutions, we processed sequencing data by reading the Pysamstats result. At each position, we extracted the reference base and the observed base counts (A, C, G, T). Positions where the base count was at least 20 were considered valid. For these valid positions, the reference base was compared to the observed bases. Then, we tracked how often each reference base was substituted by another base. Additionally, we computed the overall fraction of each base substitution by summing the base counts for each reference nucleotide.

To determine the number of genes observed in each of the datasets, we intersected each read’s aligned coordinates with the gencode.v43 gene body coordinates. We did not restrict the number of overlapping nucleotides between the read and the gene body to be counted as an observation for a gene. A gene was considered observed in a dataset if there were at least 10 supporting reads.

m^6^A calling was performed with both Dorado and m6anet. Once called, each dataset was filtered for a minimum coverage of 20 reads aligned at a given site. m6anet and Modkit were called on the modified BAM files using:

~~~
m6anet dataprep --eventalign
m6anet inference --pretrained_model HEK293T_RNA004 --num_iterations 20
modkit pileup --motif A 0
~~~

For RNA002 data, polyA tail lengths were estimated using nanopolish.

~~~
nanopolish polya
~~~

For RNA004 polyA, tail lengths were estimated using Dorado.

~~~
Dorado basecalled --estimate-poly(A)
~~~

We grouped aligned reads into mitochondrial-sourced transcripts and genomic transcripts based on the aligned chromosome for each read. Statistical testing between the discrete non-parametric populations was performed with a Wilcoxon rank-sum test.

### Analysis of data for synthetic oligonucleotides

To compare Dorado modification likelihood (ML) values for synthetic oligonucleotides and their IVT pair, we captured the read position, reference position, base call, and ML tags for each read. The ML tag is representative of the model’s confidence that a given base is modified. This allowed us to compare the model’s performance at the known modified sites and two matching bases around it (uridine for Ψ or adenosine for m^6^A) in both the modified and IVT negative conditions.

To evaluate the performance of Dorado on modified and canonical bases, we made confusion matrices for the modified positions and IVT pairs. This shows a detailed summary of Dorado’s performance by comparing the predicted condition with the true condition. True Positives (TP): where the modified base was correctly identified as modified. True Negatives (TN): where the canonical base was correctly identified as canonical. False Positives (FP): where the canonical base was incorrectly identified as modified. False Negatives (FN): where the modified base was incorrectly identified as canonical. The summary statistics of model performance were computed (**Table 1**) using the following formulas:

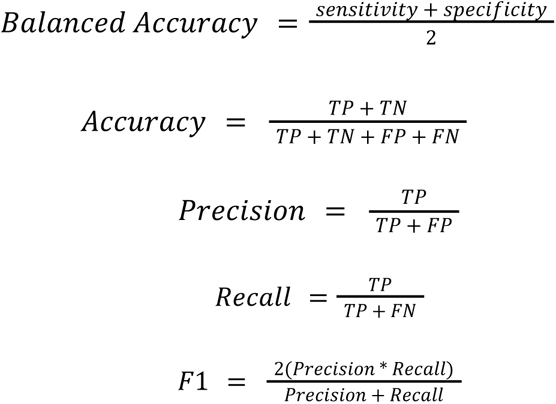

To identify putative modification sites from the GM12878 data, we selected the filtering criteria of a minimum of 20 reads with reference matching nucleotides aligned at the position (not mismatch, insertion, or deletion) with a minimum of 20% predicted modification occupancy. These criteria were selected based on the false positive rate observed using the synthetic oligonucleotide sequencing results. Assuming the highest false positive rate of 0.0219 as the success parameter for a binomial distribution, our filtering criteria give a probability of a false positive putative modification site as 0.00084 or less than 1 in 1,000 sites. We used this criterion to limit false-positive sites.

To identify isoform-specific modifications, we used Isoquant^28^, which produced a per-read isoform classification.

~~~
isoquant.py --data_type nanopore
~~~

Using the Modkit extract calls feature intersected with the isoform classification, we made an isoform-specific modification pileup. Isoform positions were transformed to genomic positions and isoform-specific modification differences were calculated.

~~~
modkit extract calls
~~~

### Reagents

RPMI medium (Invitrogen cat no. 21870076), FBS (Lifetech cat no. 12483020), L-Glutamax (Lifetech cat no. 35050061), TRI-Reagent (Invitrogen AM9738), NEXTflex Poly(A) Beads (BIOO Scientific cat. no. NOVA-512980), Direct RNA Sequencing Kit (Oxford Nanopore Technologies SQK-RNA004), T4 RNA Ligase 2 (NEB #M0239), 5X Quick Ligase Buffer (NEB #B6058), 2x Long AMP Taq (NEB #M0323), HiScribe NEB T7 High Yield RNA synthesis kit (#E2040), DNase I (NEB #M03030), SPRI beads (Beckman Coulter #B23318)

Biological Resources

GM12878 cell culture (Coriell Institute)

## Data and Code Availability

All the scripts used in this study are available in the GitHub repository (https://github.com/genometechlab/RNA002_vs_RNA004). GM12878 RNA002 sequence level data are publicly available from AWS (https://github.com/nanopore-wgs-consortium/NA12878/blob/master/RNA.md), raw signal files are available on request from the authors. GM12878 RNA004 sequence level data will be publicly available through the European Nucleotide Archive (https://www.ebi.ac.uk/ena/). Signal files are available on request from the authors.

## Acknowledgements

The authors are grateful for support from Hugh Olsen (UCSC) for advice. Elizabeth Snell, Adrien Leger, Rosemary Dokos, and Lakmal Jayasinghe (ONT) for helpful advice during RNA004 chemistry early access and the project. Arthur Rand, Adrien Leger, and Marcus Stoiber (ONT) provided advice for Dorado modification calling and Modkit. The authors also acknowledge Robin Abu-Shumays and David Garcia for helpful comments. The authors would like to acknowledge Joseph Roffi and the ISEC-Ops team for maintaining the facilities and promoting a healthy working environment.

## Author Contributions

N.G.E, A.J.S, and S.A contributed to experimental work, data analysis, writing, and editing. T.T performed experimental work and editing. M.J contributed to conceptualization, experimental work, data analysis, writing, and editing.

## Supplemental Data

Supplemental Info

## References

1. Workman, R. E. et al. Nanopore native RNA sequencing of a human poly (A) transcriptome. Nat. Methods 16, 1297–1305 (2019).

2. Garalde, D. R. et al. Highly parallel direct RNA sequencing on an array of nanopores. Nature Methods 15, 201–206 (2018).

3. Liu, H. et al. Accurate detection of m6A RNA modifications in native RNA sequences. Nature Communications 10, 1–9 (2019).

4. Leger, A. et al. RNA modifications detection by comparative Nanopore direct RNA sequencing. Nat Commun 12, 7198 (2021).

5. Huang, S. et al. Interferon inducible pseudouridine modification in human mRNA by quantitative nanopore profiling. Genome Biology 22, 1–14 (2021).

6. Tavakoli, S. et al. Semi-quantitative detection of pseudouridine modifications and type I/II hypermodifications in human mRNAs using direct long-read sequencing. Nature Communications 14, 1–12 (2023).

7. Chen, H. et al. Cross-talk of four types of RNA modification writers defines tumor microenvironment and pharmacogenomic landscape in colorectal cancer. Molecular Cancer 20, 1–21 (2021).

8. Shi, H., Chai, P., Jia, R. & Fan, X. Novel insight into the regulatory roles of diverse RNA modifications: Re-defining the bridge between transcription and translation. Molecular Cancer 19, 1–17 (2020).

9. Wang, X. et al. N6-methyladenosine-dependent regulation of messenger RNA stability. Nature 505, 117–120 (2013).

10. Cui, L. et al. RNA modifications: importance in immune cell biology and related diseases. Signal Transduction and Targeted Therapy 7, 1–26 (2022).

11. Yang, L. et al. Emerging role of RNA modification and long noncoding RNA interaction in cancer. Cancer Gene Therapy 31, 816–830 (2024).

12. Hendra, C. et al. Detection of m6A from direct RNA sequencing using a multiple instance learning framework. Nature Methods 19, 1590–1598 (2022).

13. Hewel, C., et al. Direct RNA sequencing enables improved transcriptome assessment and tracking of RNA modifications for medical applications. bioRxiv 2024.07.25.605188 (2025) doi:10.1101/2024.07.25.605188.

14. Modifications - Dorado Documentation. https://dorado-docs.readthedocs.io/en/latest/basecaller/mods/.

15. Nanopore sequencing accuracy. Oxford Nanopore Technologies https://nanoporetech.com/platform/accuracy.

16. Mulroney, L., Birney, E., Leonardi, T. & Nicassio, F. Using Nanocompore to Identify RNA Modifications from Direct RNA Nanopore Sequencing Data. Curr Protoc 3, e683 (2023).

17. Thomas, N. K. et al. Direct Nanopore Sequencing of Individual Full Length tRNA Strands. ACS Nano 15, 16642–16653 (2021).

18. Fanari, O. et al. Probing enzyme-dependent pseudouridylation using direct RNA sequencing to assess epitranscriptome plasticity in a neuronal cell line. Cell Syst 101238 (2025).

19. Shaw, E. A. et al. Combining Nanopore direct RNA sequencing with genetics and mass spectrometry for analysis of T-loop base modifications across 42 yeast tRNA isoacceptors. Nucleic Acids Res 52, 12074–12092 (2024).

20. Wang, Z. et al. Training data diversity enhances the basecalling of novel RNA modification-induced nanopore sequencing readouts. Nat Commun 16, 679 (2025).

21. Tzadikario, T. et al. Genomic in vitro transcription and Nanopore direct RNA sequencing of a human B-Lymphocyte cell line. bioRxiv (2025) doi:10.1101/2025.04.25.650674.

22. Homo sapiens genome assembly GRCh38.p14. NCBI https://www.ncbi.nlm.nih.gov/datasets/genome/GCF_000001405.40/.

23. Smith, A. M., Jain, M., Mulroney, L., Garalde, D. R. & Akeson, M. Reading canonical and modified nucleobases in 16S ribosomal RNA using nanopore native RNA sequencing. PLoS One 14, e0216709 (2019).

24. Mao, Y. et al. m6A in mRNA coding regions promotes translation via the RNA helicase-containing YTHDC2. Nature Communications 10, 1–11 (2019).

25. Roundtree, I. A., Evans, M. E., Pan, T. & He, C. Dynamic RNA Modifications in Gene Expression Regulation. Cell 169, 1187–1200 (2017).

26. Fu, Y., Dominissini, D., Rechavi, G. & He, C. Gene expression regulation mediated through reversible m^6^A RNA methylation. Nat Rev Genet 15, 293–306 (2014).

27. Wang, X. et al. N(6)-methyladenosine Modulates Messenger RNA Translation Efficiency. Cell 161, 1388–1399 (2015).

28. Prjibelski, A. D. et al. Accurate isoform discovery with IsoQuant using long reads. Nat Biotechnol 41, 915–918 (2023).

29. Akbari, V. et al. Megabase-scale methylation phasing using nanopore long reads and NanoMethPhase. Genome Biology 22, 1–21 (2021).

30. Cretu Stancu, M., et al. Mapping and phasing of structural variation in patient genomes using nanopore sequencing. Nature Communications 8, 1–13 (2017).

31. Modifications - Dorado Documentation. https://dorado-docs.readthedocs.io/en/latest/basecaller/mods/.

32. Li, H. Minimap2: pairwise alignment for nucleotide sequences. Bioinformatics 34, 3094–3100 (2018).

33. GitHub - alimanfoo/pysamstats: A fast Python and command-line utility for extracting simple statistics against genome positions based on sequence alignments from a SAM or BAM file. GitHub https://github.com/alimanfoo/pysamstats.

34. Zheng, Z. et al. Symphonizing pileup and full-alignment for deep learning-based long-read variant calling. Nature Computational Science 2, 797–803 (2022).

35. GitHub - nanoporetech/modkit: A bioinformatics tool for working with modified bases. GitHub https://github.com/nanoporetech/modkit.

36. GENCODE - Human Release 43. https://www.gencodegenes.org/human/release_43.html.

37. Loman, N. J., Quick, J. & Simpson, J. T. A complete bacterial genome assembled de novo using only nanopore sequencing data. Nat Methods 12, 733–735 (2015).

38. Gamaarachchi, H. et al. GPU accelerated adaptive banded event alignment for rapid comparative nanopore signal analysis. BMC Bioinformatics 21, 1–13 (2020).

39. Robinson, J. T. et al. Integrative genomics viewer. Nature Biotechnology 29, 24–26 (2011).

40. Direct RNA sequencing (SQK-RNA004). Oxford Nanopore Technologies https://nanoporetech.com/document/direct-rna-sequencing-sqk-rna004 (2023).

